# Gene coexpression analysis identifies genes associated with chlorophyll content and relative water content in pearl millet

**DOI:** 10.1101/2023.03.16.532967

**Authors:** Harshraj Shinde, Ambika Dudhate, Atul Sathe, Neha Paserkar, Sopan G. Wagh, Ulhas S. Kadam

## Abstract

Pearl millet is a significant crop tolerant to abiotic stresses and is a staple food of arid regions. However, its underlying mechanisms of stress tolerance is not fully understood. Plant survival is regulated by the ability to perceive a stress signal and induce appropriate physiological changes. Here, we screened for genes regulating physiological changes such as chlorophyll content (CC) and relative water content (RWC) in response to abiotic stress. Using ‘weighted gene co-expression network analysis’ (WGCNA) and clustering associated changes in physiological traits, *i*.*e*., CC and RWC with gene expression. A group of genes showing correlation with traits was identified as modules, and different color names were used to denote a particular module. In WGCNA, the darkgreen module (7082 genes) showed a significant positive correlation with CC, and the black (1393 genes) module was negatively correlated with CC and RWC. Analysis of the module positively correlated with CC highlighted ribosome synthesis and plant hormone signaling as the most significant pathways. *Potassium transporter 8* and *monothiol glutaredoxin* were reported as the topmost hub genes in the darkgreen module. In *Clust* analysis, 2987 genes were found to display a correlation with increasing CC and RWC. Further, the pathway analysis of these clusters identified ribosome and thermogenesis as positive regulators of RWC and CC, respectively. Our study provides novel insights into the molecular mechanisms regulating CC and RWC in pearl millet.

## 1. Introduction

The global human population is predicted to reach 9 billion by 2050, and at the same time, global temperature is also expected to be increased by 1 to 4 °C (1). Given those predictions, there is a need to study abiotic stress-tolerant staple food crops such as pearl millet to ensure food security. Pearl millet is an agronomically robust crop with an excellent nutritional profile and exceptional abiotic stress tolerance capacity (2). Pearl millet can flower at 42 °C, grow at 250 mM of NaCl, and produces grain at mean precipitation of as low as 250 mm (3). Despite these characteristics, pearl millet is often considered an orphan crop as it lags other staple crops in research and development (4–6). Since its genome sequencing in 2017 (7), substantial research on pearl millet has been carried out.

Several research efforts have focused on understanding pearl millet’s physiological and molecular responses to abiotic stresses (8). A previous physiological study by Shinde et al. revealed that the pearl millet tolerant line ICMB 01222 had a higher growth rate and accumulated higher sugar in leaves under salinity stress than the susceptible line (9). A study using drought-responsive pearl millet lines exhibited that drought-susceptible pearl millet line ICMB 863 showed a greater reduction in CC or greenness than drought-tolerant line ICMB 843. This study also reported photosynthesis, plant hormone signal transduction, and mitogen-activated kinase signaling as a drought-responsive pathway (10). The function of several abiotic stress-responsive genes and small RNAs has been studied in pearl millet (11–15) However, knowledge of molecular mechanisms regulating physiological responses is equally critical and important for ensuring the quality and yield of plants. Technological advancements in sequencing have created unprecedented opportunities to study those molecular mechanisms.

WGCNA is a widely used free-scale coexpression network analysis technique that constructs the modules containing genes, which shows a correlation with complex traits (16). WGCNA has been widely used to study complex plant traits and their correlation with gene expression data (17). In rice, WGCNA analysis identified CaM (Calmodulin), DUF630/632 (domain of unknown function 630/632), CHL27 (Chlamydomonas 27), and LEA4-5 (Late Embryogenesis Abundant 4-5) as a hub gene (central regulators) of salt stress-related traits (18). These hub genes will be useful for developing salt-tolerant rice genotypes.

In the current study, stress-induced physiological changes in CC and leaf water content were associated with gene expression data. We performed the WGCNA analysis (16) to generate two coexpression networks incorporating these physiological variables to determine modules of genes whose expression was regulated in concert with the physiological changes. Two interesting coexpression modules were identified, which show a positive and negative correlation with CC and RWC. For comparison, we performed the co-expression analysis using *Clust* (19). Metabolic pathway analyses of these modules and clusters were also performed. In the future, our findings might help researchers to develop strategies for the biotechnological improvement of stress tolerance in pearl millet.

## 2. Results

### 2.1. Transcript quantification

After quality assessment and filtering, such as removing adaptor sequences and discarding low-quality reads, 89.2% clean reads were generated on average. These clean reads were processed for transcriptome analysis. The clean reads mapped to the pearl millet genome. The average percentage of mappable reads per sample was 37.27%. Since transcripts Per Million (TPM) is the most used normalization method, which normalizes all the reads within a sample, the sum of all the reads would be exactly 1,000,000. The TPM for all 38,396 pearl millet genes was estimated, and the average TPM per sample was 26.2. Of 38,396 genes, 19002 had ≥1 TPM value (Supplementary file S2).

### 2.2. Weighted gene coexpression network analysis (WGCNA)

The WGCNA analysis constructed 18 modules (clusters of coexpressed genes) differentiated by color. The WGCNA also allowed us to associate the correlation between the genes of each module and traits. Figure 1 shows the association between modules and characteristics, typically representing Pearson’s correlation coefficients measured between every module and physiological characteristic. Because of the vast amount of data, we decided to focus on two modules, *i*.*e*., darkgreen and black. The darkgreen module (7082 genes) shows a positive correlation with CC (*R=0*.*71, p < 0*.*002*). In contrast, the black module (genes) has a negative correlation with both traits (with CC, *R=-0*.*51, p < 0*.*04* and with RWC, *R=-0*.*56, p <0*.*02*). Data for all genes, their respective modules (18 eigengene modules), and correlation values are given in Supplementary File S3. The hub genes within the dark magenta module related to CC were detected. The top five hub genes were the *potassium transporter 8, nonothiol glutaredoxin, chaperonin chloroplastic, oxoglutarate-dependent dioxygenase*, and *uncharacterized protein* (Figure 2).

**Figure 1.**
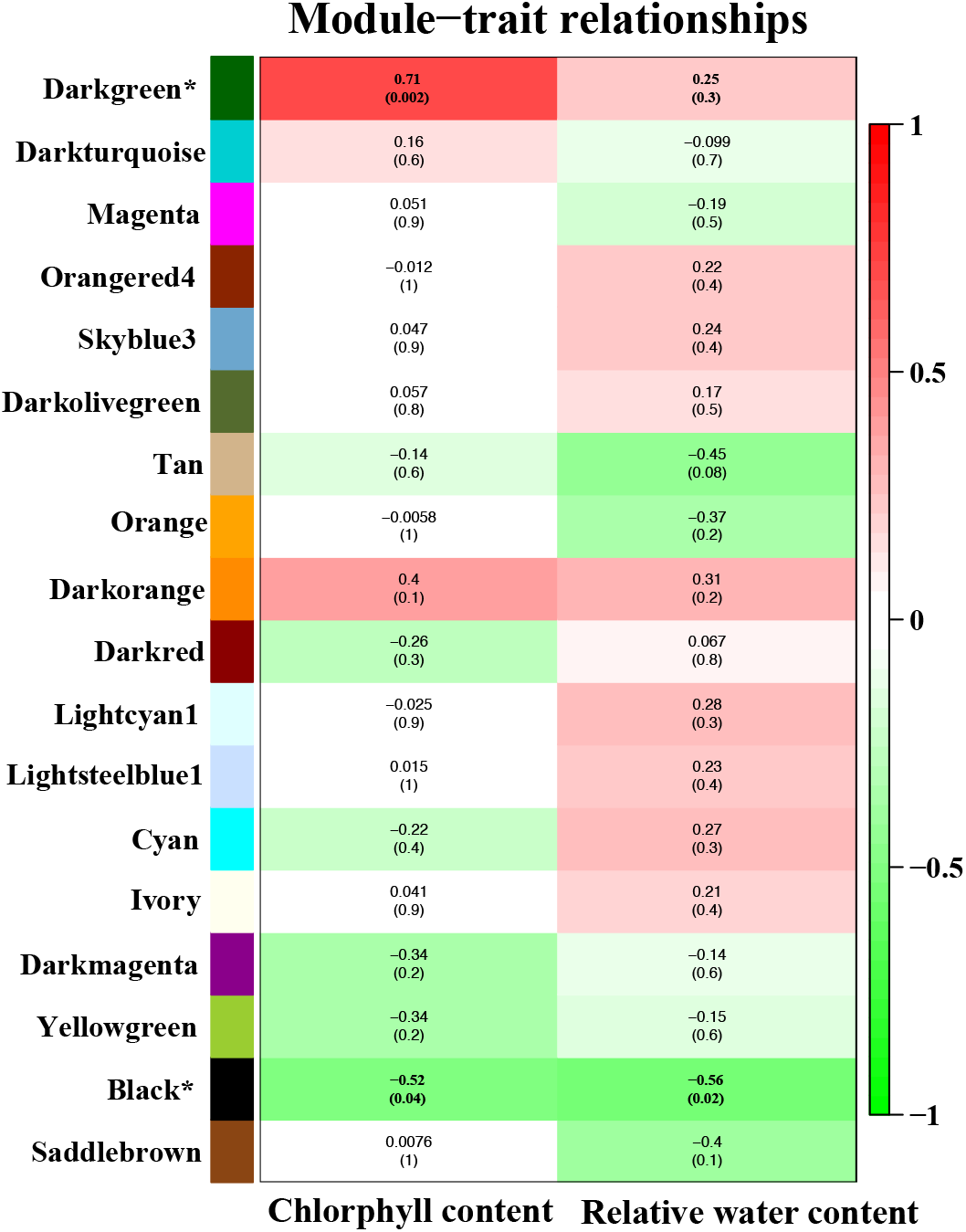
Correlation between modules (group of genes) and traits [CC and RWC]. Module names are displayed on the left, and the correlation coefficients are shown at the top of each row. The p-values for each module are displayed at the bottom of each row within parentheses. The rows are colored based on the correlation of the module with the trait: Red for positive and green for the negative correlation.

**Figure 2.**
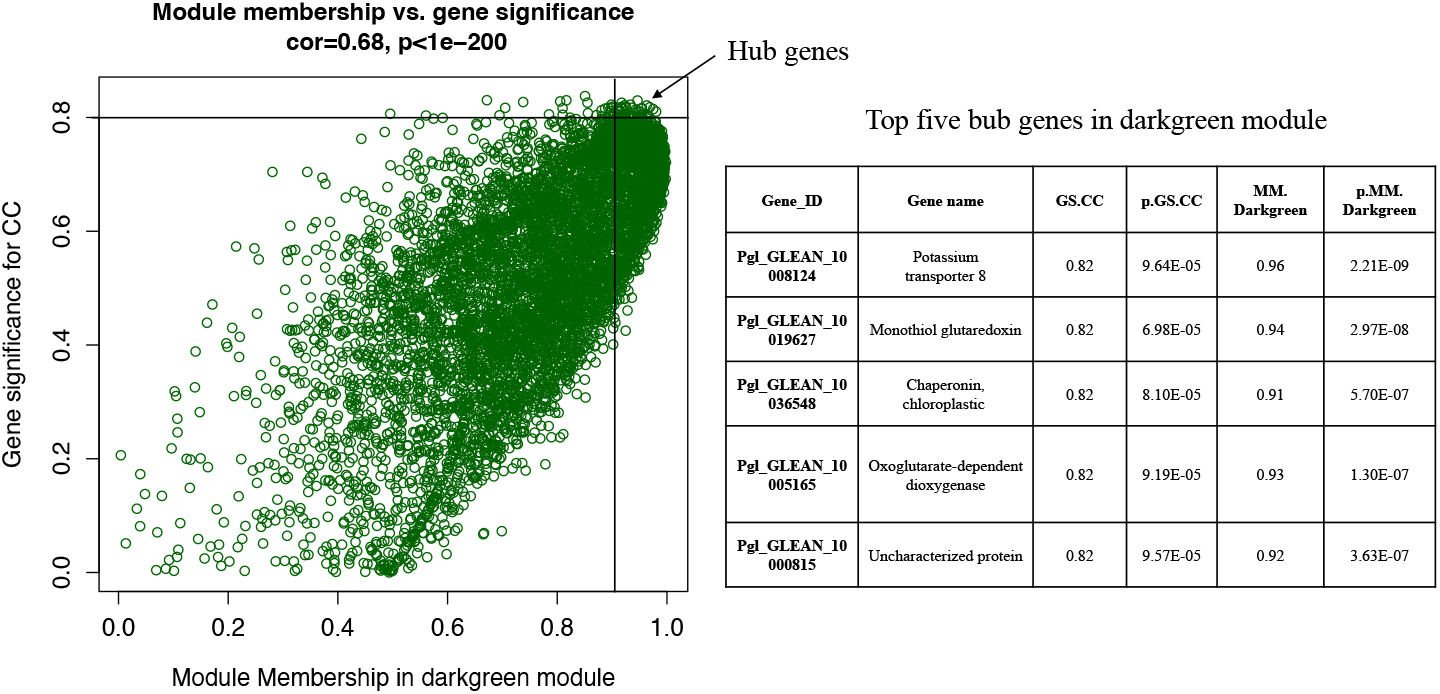
Hub genes are determined by module membership (MM) and gene significance (GS). GS represents the correlation between a gene and a trait. The MM represents the correlation between an individual gene and the module eigengene. MM and GS values were plotted on the x and y-axis. In the darkgreen module, hub genes were identified according to a GS >0.8 and an MM >0.9. The table represents the top five hub genes.

### 2.3. Gene clustering analysis

Cluster analysis results were divided into two categories: CC-related genes and RWC-related genes. For CC, 12 clusters were formed (C0 to C11). The largest cluster was C0 (4699 genes), while the smallest cluster was C11 (59 genes) (Supplementary Figure 1). Among these clusters, C8 (603 genes) and C2 (835 genes) were selected for pathway analysis as they exhibited either positively or negatively coexpressed linear expression patterns with increasing chlorophyll content. For CC, the list of genes in each cluster is given in supplementary file S4.

Similarly, for RWC, in total, 11 clusters were formed (C0 to C11). The largest cluster was C0 (3090 genes), while the smallest cluster was C3 (66 genes) (Supplementary Figure 2). From these 11 clusters, C11 (828 genes) and C5 (721 genes) were selected for further analysis as they exhibited linear expression patterns either positively or negatively coe-pressed with increasing RWC. For RWC, the list of genes present in each cluster is given in supplementary file S5.

### 2.4. Metabolic pathway analysis

Pathway analysis of genes in the ‘darkgreen’ module (positively correlated with chlorophyll content) showed them to be enriched in metabolic pathways such as the ribosome, plant hormone signal transduction, starch and sugar metabolism, photosynthesis, and MAPK signaling. Ribosome synthesis was the most enriched pathway with 48 genes associated (Figure 3).

**Figure 3.**
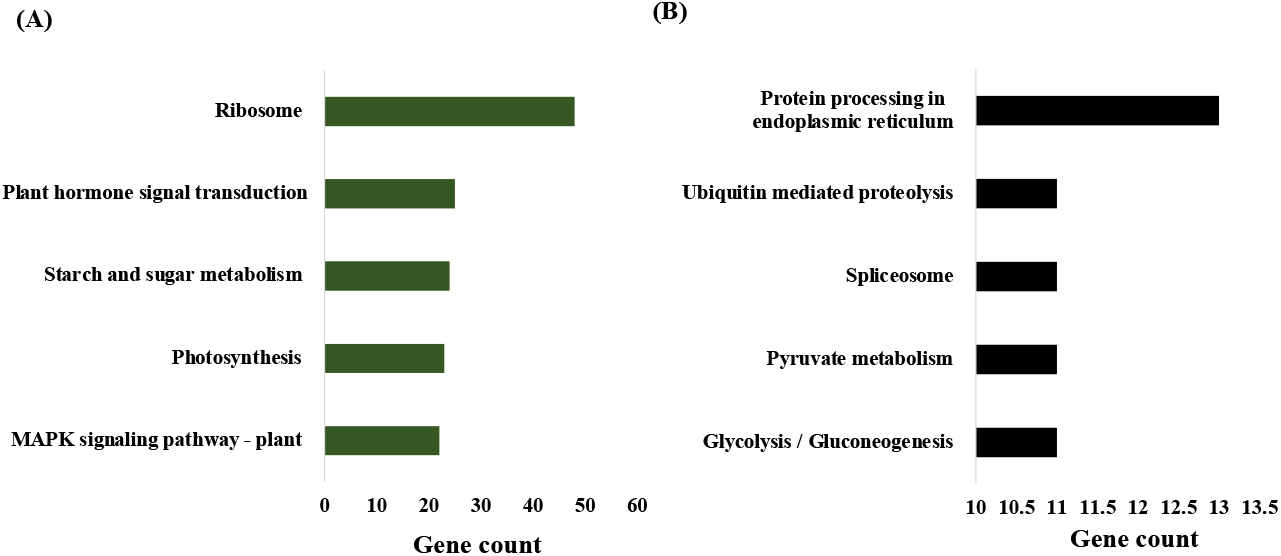
Metabolic pathway analysis of significant modules. (A) Pathways of darkgreen module (positively correlated) (B). Pathways of the black module (negatively correlated).

In linearly coexpressed clusters, ribosome, protein processing in the endoplasmic reticulum, plant-pathogen interaction, plant hormone signal transduction, and phenylpropanoid biosynthesis were designed as a positive regulator of RWC. Endocytosis, starch and sucrose metabolism, phenylalanine, tyrosine and tryptophan biosynthesis, mRNA surveillance pathway, and RNA degradation were negative regulators of RWC. (Figure 4). In the case of CC clusters, thermogenesis, aminoacyl-tRNA biosynthesis, glycolysis/gluconeogenesis, an amino sugar, nucleotide sugar metabolism, and pyruvate metabolism as positive regulators of chlorophyll content. Whereas Plant hormone signal transduction, Protein processing in the endoplasmic reticulum, Ribosome biogenesis in eukaryotes, Nucleocytoplasmic transport, and phenylpropanoid biosynthesis as a negative regulator of CC (Figure 5).

**Figure 4.**
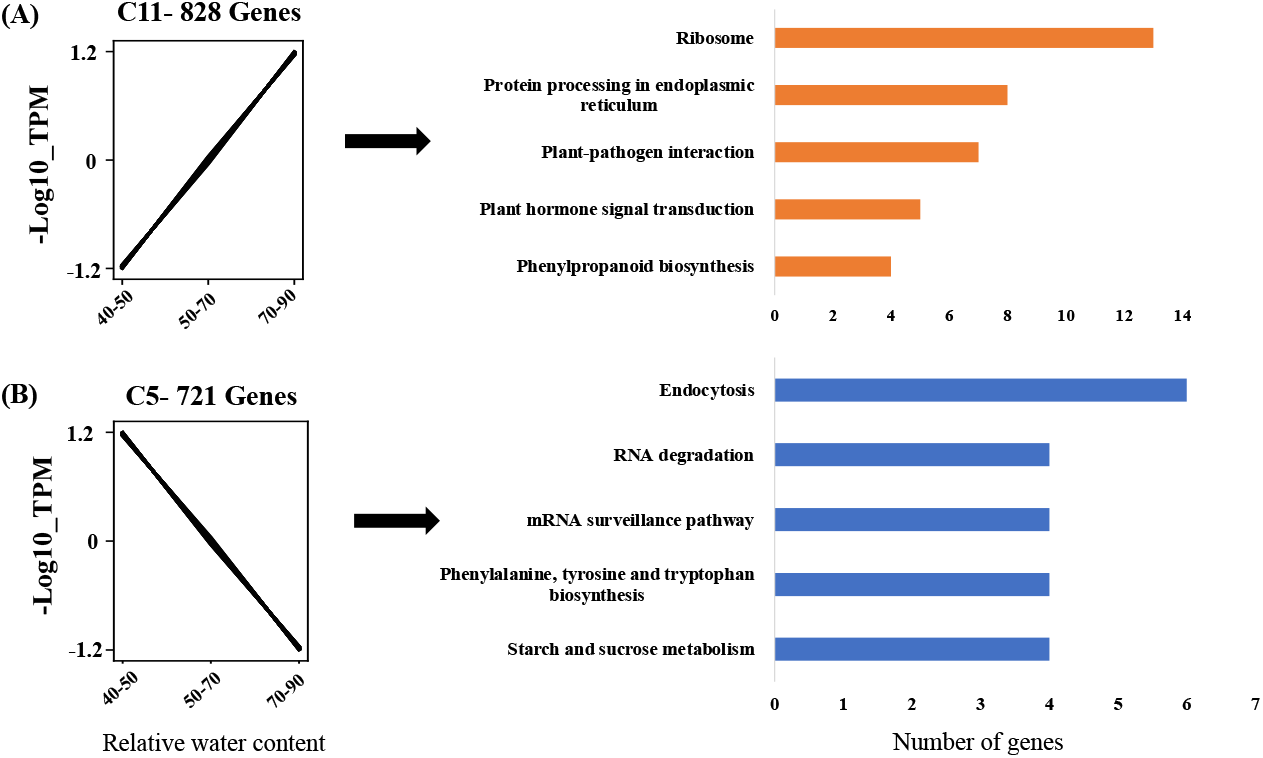
Clusters of coexpressed genes related to RWC (RWC) elucidated by *Clust*. (A) Gene cluster showing positive correlation with RWC, n = 828, and its pathways. (B) Gene cluster showing negative correlation with RWC, n = 721, and its pathways.

**Figure 5.**
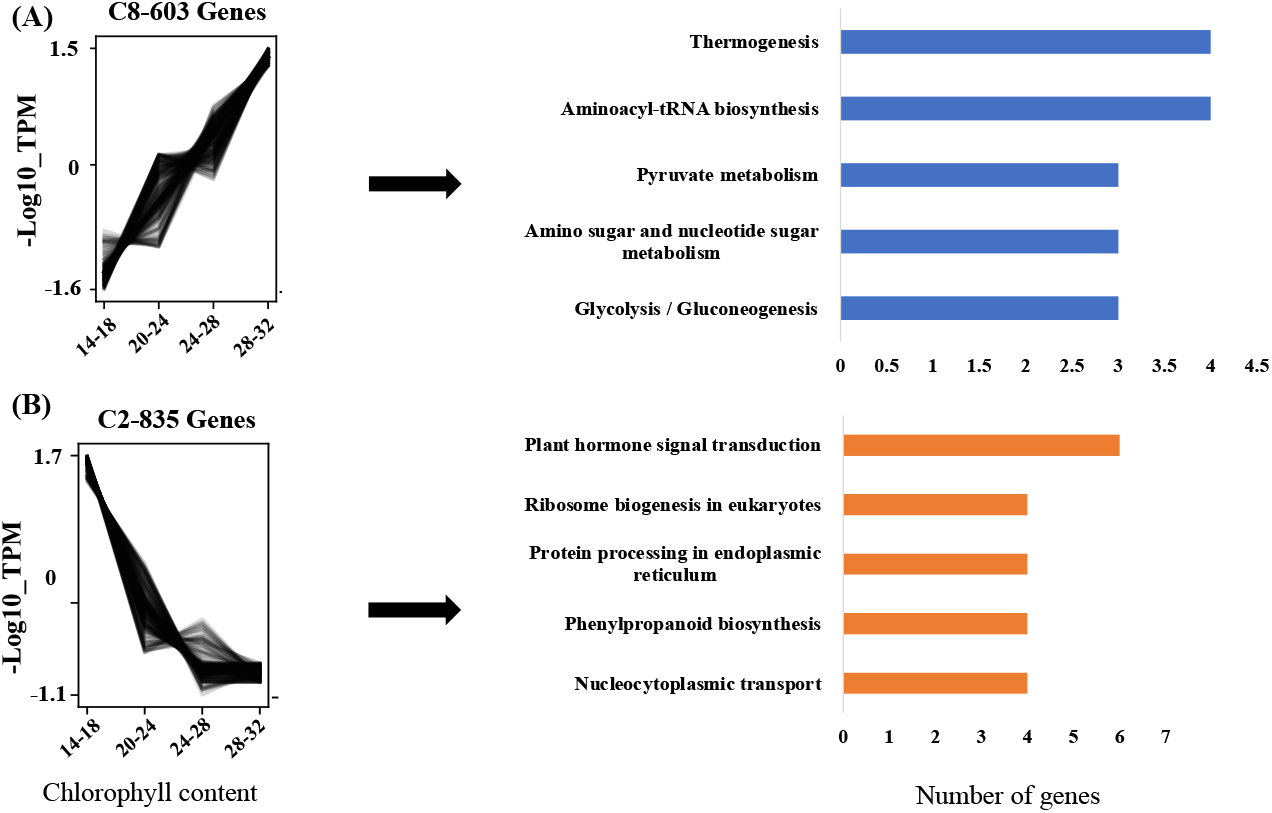
Clusters of coexpressed genes related to CC (CC) elucidated by *Clust*. (A) Gene cluster showing positive correlation with CC, n = 603; and its pathways. (B) Gene cluster showing negative correlation with CC, n = 835, and its pathways.

Our WGCNA and *Clust* analysis results aligned well with each other as we observed that ribosome and plant hormone signal transduction are positive regulators of RWC in both analyses.

## 4. Discussion

Pearl millet is a staple crop of arid and semi-arid tropics and is known for its unique ability to tolerate abiotic stress. Various studies used a genomic approach to identify genes and pathways related to abiotic stress in pearl millet (20–25). But none of the studies performed a coexpression analysis to discover genes that regulate abiotic stress-related traits. CC and RWC are abiotic stress-related traits and change rapidly during abiotic stresses.

Integrating these dynamic changes in physiology with the transcriptome data using WGCNA and *Clust* approach uncovered genes regulating ribosome, plant hormone signal transduction, protein processing, thermogenesis, and endocytosis. Among those pathways, the ribosome is the most important pathway as it is found to be a positive regulator of RWC in both WGCNA and *Clust* analysis. The ribosomes are responsible for synthesizing proteins. (26) The functioning of the ribosome during stress is crucial for the timely synthesis of stress-responsive proteins. (27,28)Different stress and defense-related pathways get deactivated upon ribosomal impairment (29). This crosstalk between ribosome and stress-responsive protein serves as a novel approach to studying stress-tolerant plants. Plant hormone signaling and protein processing were other pathways reported in WGCNA modules and clusters. As plant hormones are early responders to stress stimulus (30), we suggest early sensing of stress by pearl millet at the molecular level.

Endocytosis, protein processing in the endoplasmic reticulum, and plant hormone signaling were the pathway showing a negative correlation with both CC and RWC. A previous study in the *Medicago* plant has reported the relation between drought stress and endocytosis (31) where the endocytosis affects membrane lipids composition during stress. The endoplasmic reticulum is the organelle where most of the proteins of a cell are synthesized. Such proteins later play a significant role (32) in stress response and protection.

We identified hub genes in the darkgreen module, which were highly correlated with the CC trait. The topmost hub gene we found in this study is potassium transporter 8. Potassium transporters protect plant leaves from Na+ over-accumulation and salt stress during salt stress (33) Future experiments must validate the modules, clusters, metabolic pathways, and hub genes discovered in the present study.

## 4. Materials and Methods

### 4.1. RNA sequencing data pre-processing

From our previously published study (10,11), raw sequencing reads were downloaded from the NCBI-SRA database using the SRA toolkit. The quality check of raw reads was performed using FastQC (https://www.bioinformatics.babraham.ac.uk/projects/fastqc/). Adaptors and poor-quality reads were filtered using trimmomatic (34). Clean reads were then used for mapping and quantifying transcript abundance.

### 4.2. Transcript quantification

The transcriptome-wide quantification in the form of transcript per million (TPM) was performed using Salmon v1.6.0 (35). The pearl millet reference transcriptome file for mapping and quantification was obtained from the International Pearl Millet Genome Sequencing Consortium (https://cegresources.icrisat.org/data_public/PearlMillet_Genome/v1.1/). Prior to mapping and transcription quantitation, the pearl millet reference transcriptome file was indexed.

### 4.2. Weighted gene coexpression network analysis (WGCNA)

The R package, weighted gene correlation network analysis (WGCNA), was used to construct a gene coexpression network and to find coexpressed genes (13). Gene expression data and TPM of 16 plants with two different traits (CC and RWC) were used for WGCNA analysis. Trait data used in this study were also taken from our previous study. (Supplementary File S1). At first, the genes with low TPM counts and outliers were filtered. To choose modules (group of genes) associated with traits of interest, i.e., CC and RWC, Pearson correlation analysis was determined between each module’s “eigengene” and the traits. Modules with a module-trait correlation >0.5 or <-0.5 for at least one trait (P≤ 0.05) were considered significant. Gene significance (GS) was calculated for each gene as the correlation between gene expression counts and traits. Hub genes were identified by choosing genes with the highest gene significance and module membership in the significant module.

### 4.3. Gene clustering analysis

Gene expression data were clustered based on physiological measurements, i.e., CC and RWC using the Python package *Clust* v1.8.4. The plants were divided into four groups for chlorophyll content (relative greenness, SPAD reading), *i*.*e*., 10-14, 20-24, 24-28, and 28-32. The group of genes which shows increasing or decreasing trend as CC changes were selected for pathway analysis. Similarly, for RWC (relative water content in leaf, in percentage), plants were divided into three groups, *i*.*e*., 40-50, 50-70, and 70-90; and changes in gene expression were recorded.

### 4.4. Metabolic pathway analysis

We selected the significantly associated modules (R > 0.5 or <-0.5, with a p-value of 0.05) and gene clusters for the Kyoto Encyclopedia of Genes and Genomes (KEGG) pathway analysis. The KEGG pathway analyses were performed by submitting the nucleotide sequences of modules and clusters to KEGG automatic annotation KEGG Automatic Annotation Server (KAAS server located at http://www.genome.jp/kegg/kaas/) (36).

## 5. Conclusions

This study’s most relevant finding is identifying WGCNA modules [darkgreen (7082 genes) and black (1393 genes)] and clusters. These modules and clusters are enriched in the endoplasmic reticulum’s metabolic pathways like the ribosome synthesis, endocytosis, plant hormone, signal transduction, and protein processing. Based on the current finding, we postulate that during environmental stress, pearl millet can sense the stress using the hormones and transduces this signal to initiate the appropriate physiological and molecular responses and cope with stress-related perturbation. Moreover, the ribosome is involved in the synthesis of stress-responsive proteins during a stressed condition. *Potassium transporter 8* and *monothiol glutaredoxin* were the most significant hub genes in the darkgreen module. Modules, metabolic pathways, and hub genes identified in this study, provide guidelines for improving the pearl millet’s abiotic stress tolerance through future molecular breeding. The discovery of these modules, candidate hub genes can now be paired with various molecular techniques, such as gene editing, to develop climate-smart crops.

### Supplementary Materials

The following supporting information can be downloaded at

**Supplementary Figure 1.**
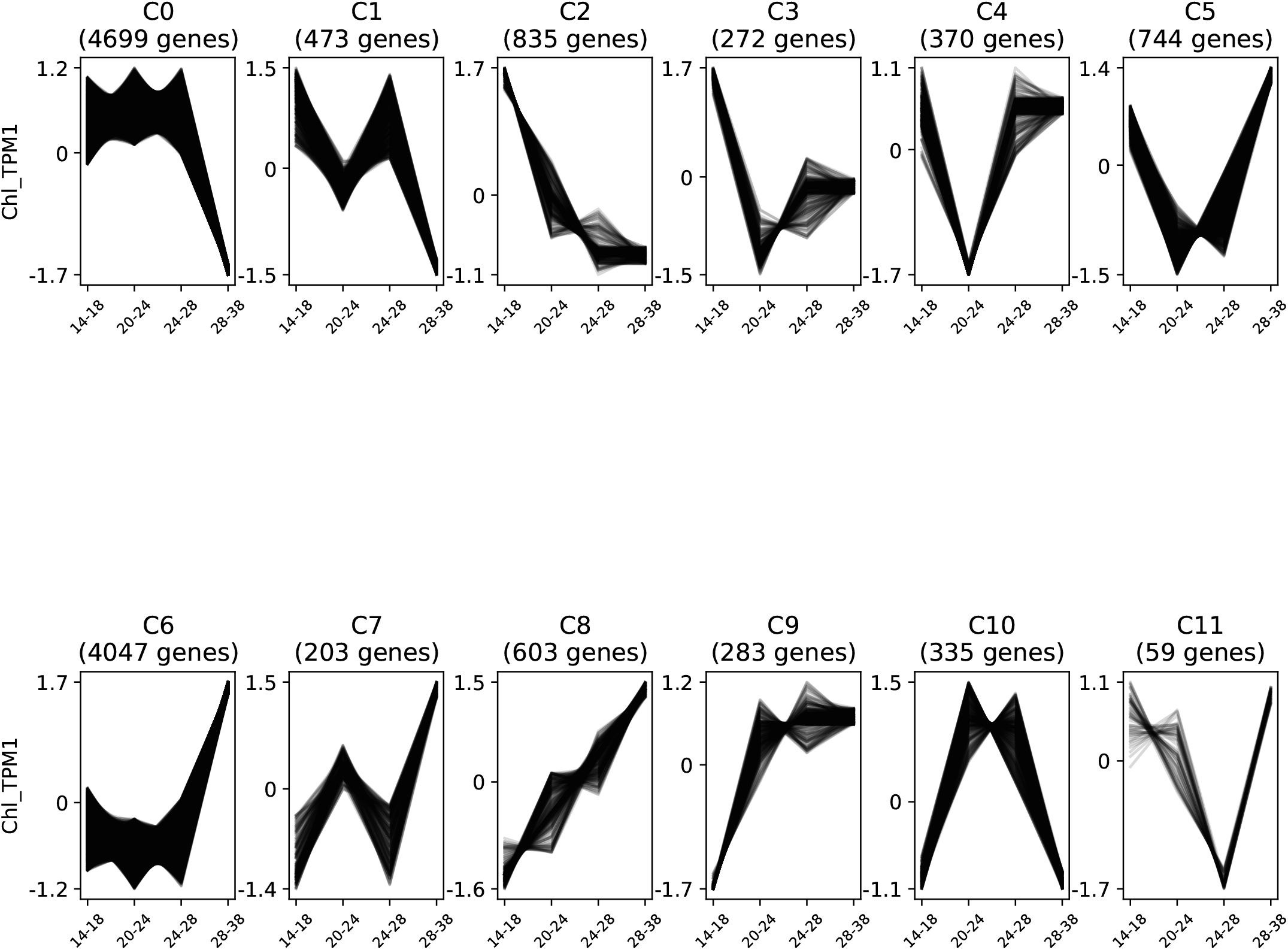
*Clust* analysis of gene clusters related to chlorophyll content. A total of 11 clusters were identified. The X-axis shows CC, and the Y-axis shows - log 10 of transcript per million values.

**Supplementary Figure 2.**
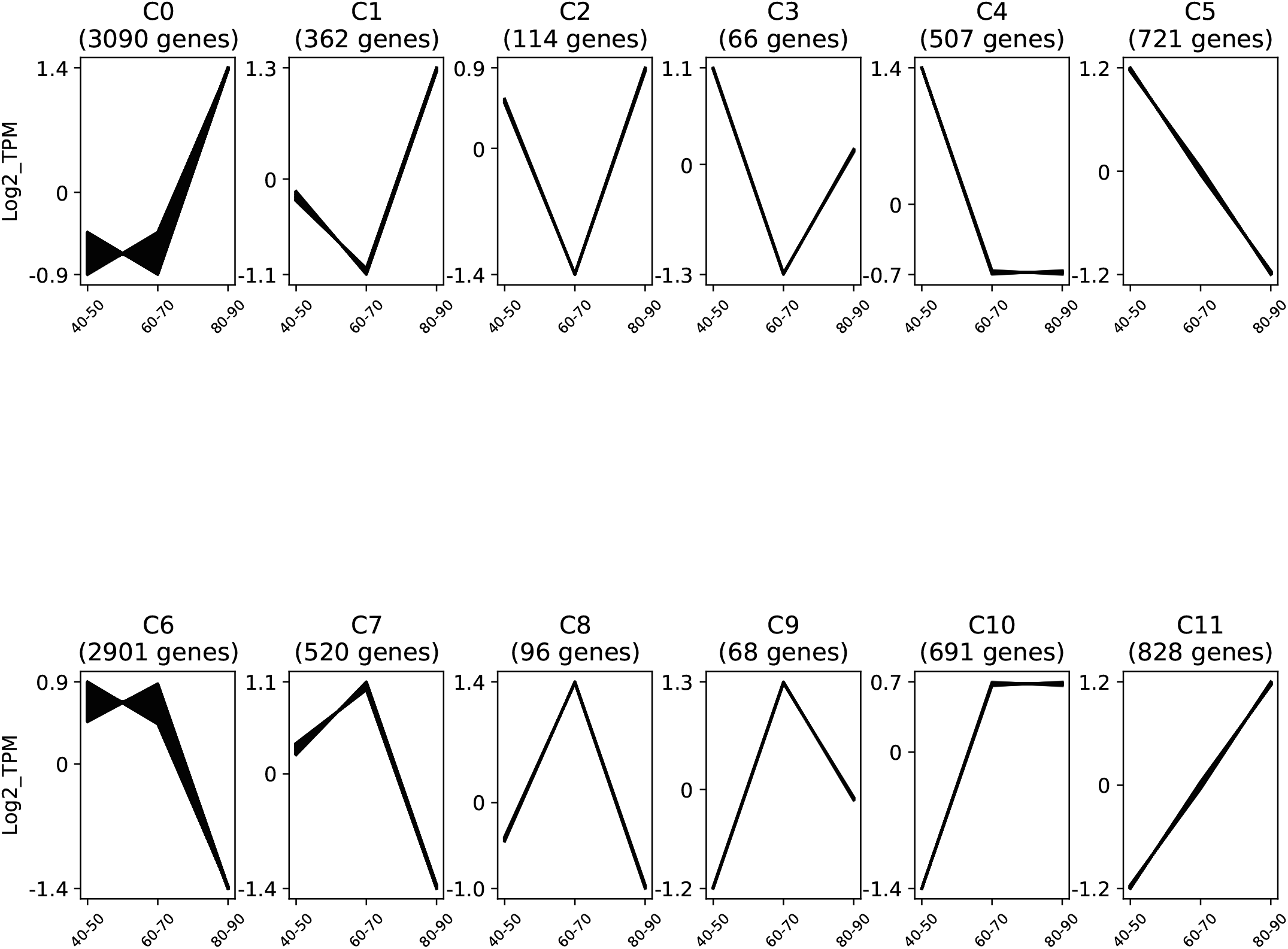
*Clust* analysis of gene clusters related to RWC. A total of 11 clusters were identified. The X-axis shows RWC, and the Y-axis shows -log_10_ of transcript per million values.

**Supplemental Table S1**. Physiological traits (CC and RWC) of 16 pearl millet plants used for gene coexpression analysis

**Supplemental Table S2**. Transcripts Per Million (TPM) values of all 16 pearl millet plants.

**Supplemental Table S3**. Data of all identified WGCNA modules and their module correlations and module membership values. (For CC and RWC)

**Supplemental Table S4**. List of pearl millet genes present in different clusters identified through *Clust*. Clusters for Chlorophyll content.

**Supplemental Table S5:** List of pearl millet genes present in different clusters identified through *Clust*. Clusters for RWC.

## Author Contributions

Conceptualization, HS. and AD.; Methodology, HS, AS, SGW and NP.; Software, AD, HS and AT.; Validation, AT, NP and HS; Formal Analysis, AT, AD and HS.; Investigation, HS and USK; Resources, HS and USK; Data Curation, HS, AD, and SGW; Writing – Original Draft Preparation, HS, AD, AT and NP.; Writing – Review & Editing, USK, SGW, and AD; Visualization, HS, SGW, and USK.; Supervision, HS and USK.; Project Administration, HS, AD and USK.; Funding Acquisition, HS and USK.

## Acknowledgment

USK was financially supported by the National Research Foundation of Korea (Grant #2022R1I1A1A01064372).

## Data Availability Statement

In this study, publicly available sequencing data sets were analyzed. Bio Project ID PRJNA419859 and SRA accession number SRP128956.

